# Cryo-EM structure of adenovirus type 3 fibre with desmoglein 2 shows a novel mode of receptor engagement

**DOI:** 10.1101/471383

**Authors:** Emilie Vassal-Stermann, Gregory Effantin, Chloe Zubieta, Wim Burmeister, Frédéric Iseni, Hongjie Wang, André Lieber, Guy Schoehn, Pascal Fender

## Abstract

Attachment of adenovirus (HAd) to host cell is a critical step of infection. This work reports the cryo-electron microscopy (cryo-EM) structure of a non-symmetrical complex smaller than 100kDa formed by the trimeric human adenovirus of type 3 fibre knob (HAd3K) and human desmoglein 2 (DSG2). The structure reveals a unique stoichiometry, shedding light to new adenovirus infection strategies and providing new insights for adenoviral vector development.

## Main Text

Desmoglein 2, a newly identified adenovirus receptor has been reported to be used by species B human adenoviruses such as HAd3, HAd7, HAd11 and HAd14 for cell infection^1^. As exemplified by HAd3 and HAd11, these viruses are used for cancer virotherapy ^2,3,4^. The structure of the extracellular region of DSG2 containing four cadherin domains EC1 to EC4, has recently been solved by crystallography^5^. The intermediate portion of the DSG2 ectodomain, consisting of EC2 and EC3, has been described as important for recognition by the HAd3 knob (HAd3K), the trimeric globular distal part of the fibre protein. HAd3K binds to DSG2 by non-classical mechanism involving mainly one receptor bound per trimeric fibre knob resulting in a sedimentation coefficient of 5.40S corresponding to a 96kDa complex^6^.

To investigate the specificity of DSG2 interaction with adenovirus, we have determined the structure of HAd3K in complex with EC2-EC3 by phase plate cryo-EM. The interaction takes place at the top of the trimeric HAd3K **(Figure 1a-b).** This observation is in stark contrast with both CD46/HAd11K and CAR/HAd12K interactions which involved the intermediate or lower part of the knob respectively^7,8^ **(Figure 1c-d).** The location of only one EC2-EC3 close to the three-fold axis is unusual as compared to CD46 and CAR for which three receptors were found at the periphery of the fibre knob. Moreover, different population of particles with either one or two EC2-EC3 modules (HAd3K / EC2-EC3 and HAd3K / (EC2-EC3)2) were identifiable **(Supplementary Figure S1.a-c)** in agreement with the 5,40S and 7,34S species previously reported^6^. The 3D structures were solved to an overall resolution of 3.5 and 3.8 Å, respectively. The resolution of both 3D maps is relatively uniform, the HAd3K and the EC2-EC3 core regions being slightly better defined than the receptor’s distal parts **(Supplementary Figure S1.d-e)**. The atomic coordinates of both HAd3K (PDB: 1H7Z) and EC2-EC3 (PDB: 5ERD) were used to fit the cryo-EM maps and refined **(Figure 1a, Supplementary Figure S1.f-k)**.

Whether one or two receptors are bound, both EC2 and EC3 domains are involved in the interaction **(Figure 2a-b, Supplementary Figure S2.a-b)**. This observation is in agreement with the biochemical data reporting that EC3 is critical for HAd3K binding while EC2 stabilizes this interaction^6^. Several residues distributed along the EC2 primary sequence were found to interact with the HAd3K monomer A, while only one region in EC3 (covering residues 313 to 321) is involved in the interaction with the groove of monomer B **(Figure 2c).** Thus, each DSG2 cadherin domains interact with one monomer **(Figure 2a-b)** as previously reported for CD46 (SCR1 and SCR2 domains) interaction with HAd11K **(Figure 1c)**. The residues at the interface of HAd3K and EC2-EC3 module have been identified **(Figure 2c-d)**. Of note, in a previous work using random mutations in the HAd3K, we have reported that single mutations of either N186D, V189G, S190P or L296R reduced DSG2 binding by more than 80% and that D261N or F265L totally abolished the binding^9^. Remarkably, all these residues were found to be involved in the interaction at the light of the present structure **(Figure 2d)**. Examples include S190/S175 or D261 forming a salt bridge with R316, stabilizing the loop conformation of HAd3, which is likely required for DSG2 binding **(Supplementary Figure S2c-d)**.

**Figure 1:**
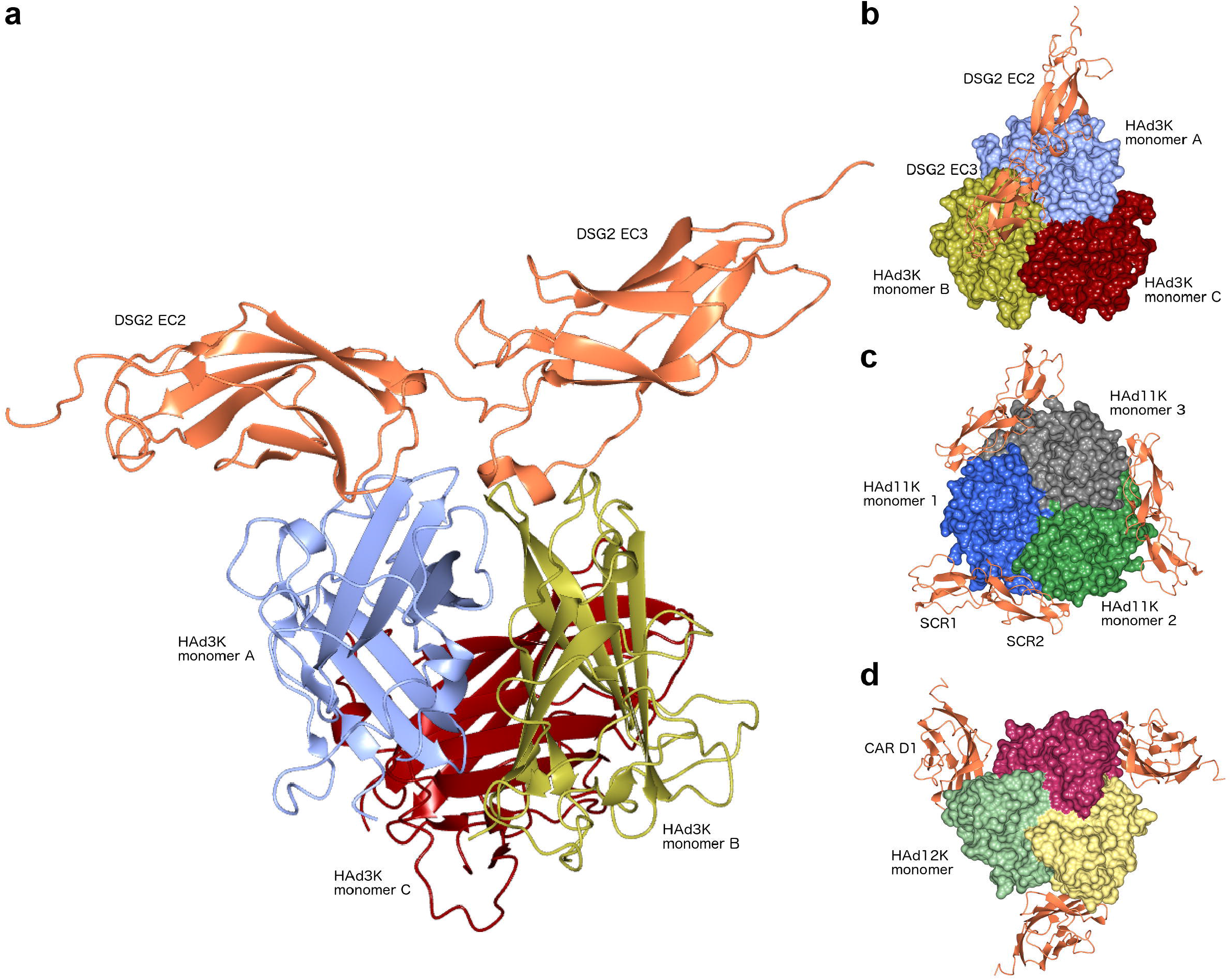
Atomic reconstruction of HAd3K/EC2-EC3 complex and comparison to other adenoviral receptors. (a) Side view of HAd3K/EC2-EC3 complex with the three HAd3K monomers colored in blue, gold and red and EC2-EC3 colored in orange. (b) Top view of the same complex. (c) Top view of HAd11 Knob structure with three CD46 SCR1&2 molecules in orange (PDB: 2039). (d) Top view of HAd12 Knob with CAR D1 domain in orange (PDB: 1KAC)

As mentioned above, beside the 5,40S complex consisting of one receptor per trimeric knob a second minor population of 7,34S (HAd3K / (EC2-EC3)_2_) has also been found. In the 7,34S complex, one of the two EC2-EC3 modules is bound similarly to the receptor of the 5.40S complex. However, the relative positions of EC2 and EC3 differ slightly in the cryo-EM maps (rmsd of 2.585 Å) likely due to the binding of the second module in the 7,34S species. The structural data provided here exclude again the three-receptor per knob *scenario*. The 1:1 (receptor:knob) binding strategy is more in agreement with both the statistical data **(Supplementary Table1)** and with our previously biochemical study^6^. Of note, multimerization of the trimeric HAd3K is necessary to cluster enough DSG2 molecules for intracellular signaling that leads to junction opening ^10,11^. Anyhow, the DSG2 interaction with adenoviruses differs from the CAR or CD46 ones.

Another notable feature of the HAdK3 / DSG2 interaction is a conformational change between the unbound and bound receptor. When superimposing the cryo-EM and crystallography structures by their EC2 domains (rmsd = 0.826 Å), it appears that the EC3 domain rotates upon interaction with HAd3K **(Figure 2e)** by ~14°. The same observation is true if the structures are superimposed by their EC3 domains (rmsd = 1.223 Å). This leads to the conclusion that the relative orientation of EC2 and EC3 is altered upon binding to HAd3K. A similar observation has been made for the binding of CD46 to HAd11K where an even more dramatic change of curvature (~60°) between CD46 SCR1 and SCR2 was observed upon binding. The HAd3 structure, in contrast does not exhibit dramatic rearrangement upon EC2-EC3 binding.

**Figure 2:**
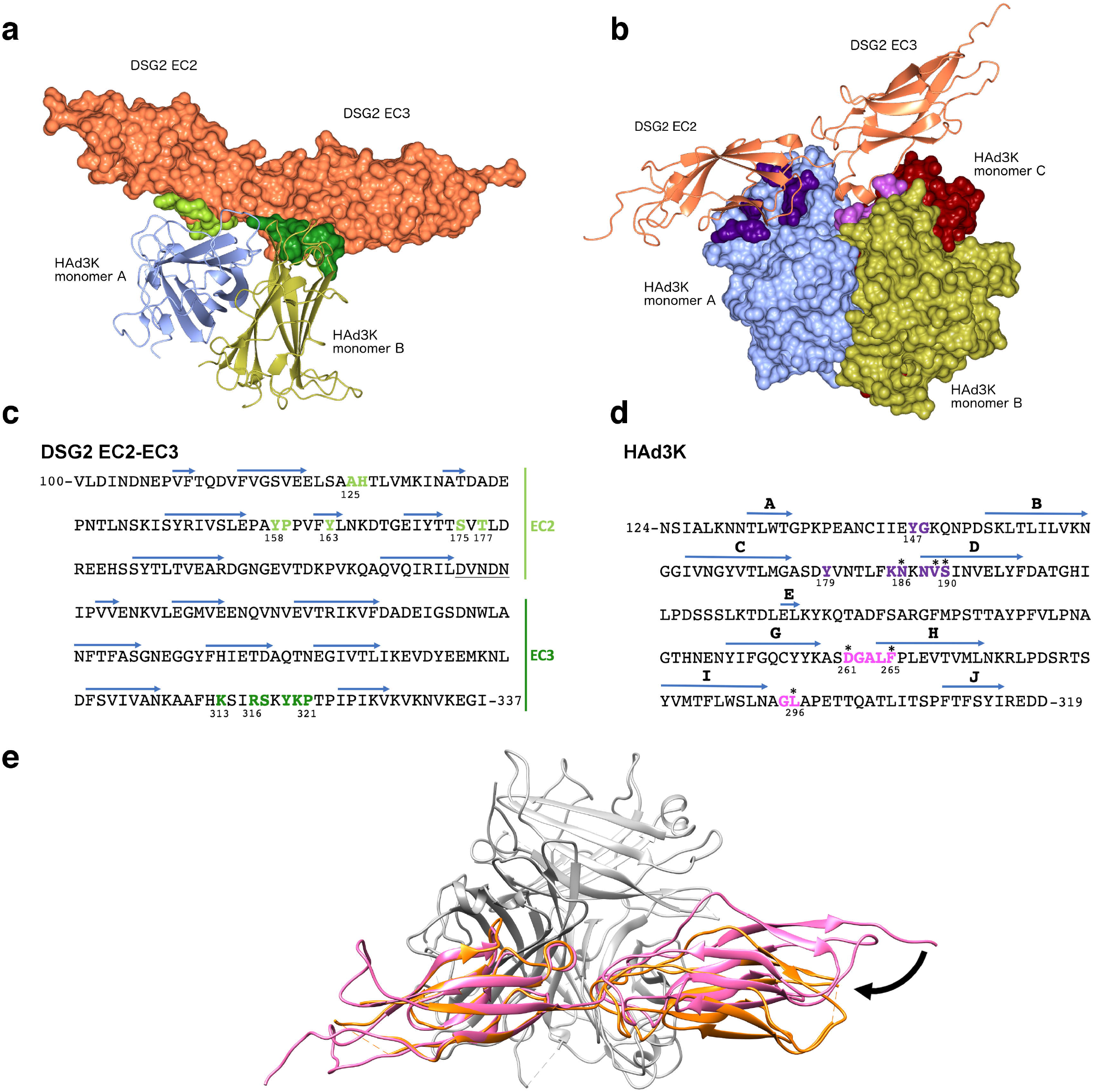
Determination of residues from HAd3K and EC2-EC3 involved in the interaction. (a) Surface of EC2-EC3 interacting with HAd3K. Aminoacids making contact with HAd3K are colored in light or dark green according to their interaction with monomer A or B, respectively. For a better view only two monomers of HAd3K are represented in cartoon. (b) Surface of HAd3K involved in receptor binding. HAd3K aminoacids involved in binding are highlighted in dark purple (EC2 contacts) and light purple (EC3 contacts). (c) EC2-EC3 primary sequence with aminoacids making contact with the knob are colored using same color code than in (a). (d) Primary sequence of HAd3K with β-strand depicted with arrows. Contact-forming residues of HAd3K are depicted with same colors as in (b). Asterisks (*) denote residues for which single mutation reduce DSG2 binding by more than 80%^9^. (e) Superimposition of free and bound EC2-EC3 (in pink and orange, respectively) with HAd3K in grey.

The data presented here reveal a new mode of interaction for DSG2 interacting adenoviruses and provides new clues for the rational design of DSG2 interacting adenoviruses, subviral particles, or fibre knobs. Such vectors are currently undergoing a rapid development in therapy with HAd3, HAd11 being used in clinical trials^2,3,4^. Beside the use of oncolytic adenoviruses, HAd3 fibre-containing molecules such as penton-dodecahedron (symmetric particle harboring 12 fibres) or junction-openers (JOs) have been reported to act as enhancer of approved treatments in cancer therapies^1,10–12^

## Author contributions

AL & PF supervised the project. EVS, GE, FI, GS designed the experiments. EVS, GE, CZ, HW and WB performed the biochemical and structural experiments. EVS, GE, GS, AL and PF wrote the manuscript.

The authors declare to have no conflict of interest or competing financial interests.

## Acknowledgments

We are grateful to the ANR for its support to the ‘Ad-Cadh’ project ANR-18-CE11-0001. We acknowledge access to the European Synchrotron Radiation Facility (ESRF), and thank CM01 scientists for their help with data collection. This work used the platforms of the Grenoble Instruct-ERIC Center (ISBG: UMS 3518 CNRS-CEA-UGA-EMBL) with support from FRISBI (ANR-10 INSB-05-02) and GRAL (ANR-10-LABX-49-01) within the Grenoble Partnership for Structural Biology (PSB). We thank Daphna Fenel and Emmanuelle Neumann for preliminar negative staining screening. The electron microscope facility is supported by the Rhône-Alpes Region, the Fondation Recherche Medicale (FRM), the Fonds Européen de Développement Régional (FEDER) and the GIS-Infrastrutures en Biologie Sante et Agronomie (IBISA).

